# Heterotrophic Carbon Fixation in a Salamander-Alga Symbiosis

**DOI:** 10.1101/2020.02.14.948299

**Authors:** John A. Burns, Ryan Kerney, Solange Duhamel

**Affiliations:** Lamont-Doherty Earth Observatory of Columbia University, Division of Biology and Paleo Environment, Palisades, NY; Bigelow Laboratory for Ocean Sciences, East Boothbay, ME; Gettysburg College, Biology, Gettysburg, PA; University of Arizona, Department of Molecular and Cellular Biology, Tucson, AZ

**Keywords:** Symbiosis, carbon fixation, photosymbiosis, mutualism, heterotroph, alga, salamander, embryo

## Abstract

The unique symbiosis between a vertebrate salamander, *Ambystoma maculatum*, and unicellular green alga, *Oophila amblystomatis*, involves multiple modes of interaction. These include an ectosymbiotic interaction where the alga colonizes the egg capsule, and an intracellular interaction where the alga enters tissues and cells of the salamander. One common interaction in mutualist photosymbioses is the transfer of photosynthate from the algal symbiont to the host animal. In the *A. maculatum*-*O. amblystomatis* interaction, there is conflicting evidence regarding whether the algae in the egg capsule transfer chemical energy captured during photosynthesis to the developing salamander embryo. In experiments where we took care to separate the carbon fixation contributions of the salamander embryo and algal symbionts, we show that inorganic carbon fixed by *A. maculatum* embryos reaches 2% of the inorganic carbon fixed by *O. amblystomatis* algae within an egg capsule after 2 hours in the light. After 2 hours in the dark, inorganic carbon fixed by *A. maculatum* embryos is 800% of the carbon fixed by *O. amblystomatis* algae within an egg capsule. Using photosynthesis inhibitors we show that *A. maculatum* embryos and O. *amblystomatis* algae compete for available inorganic carbon within the egg capsule environment. Our results confirm earlier studies suggesting a role of heterotrophic carbon fixation during vertebrate embryonic development. Our results also show that the considerable capacity of developing *A. maculatum* embryos for inorganic carbon fixation precludes our ability to distinguish any minor role of photosynthetically transferred carbon from algal symbionts to host salamanders using bicarbonate introduced to the egg system as a marker.

## Introduction

During embryonic development, egg capsules of multiple amphibian species found in the Northern Hemisphere are colonized by a green alga, *Oophila amblystomatis*. Particular attention has been given to the conspicuous association between the common spotted salamander of North America (*Ambystoma maculatum*) and its *O. amblystomatis* symbiont (Orr, 1888; Gilbert, 1942). While recent studies have focused on the unique facultative endosymbiotic association of algal cells inside the embryo host (Kerney et al., 2011, 2019; Burns et al., 2017), there is a long history of research into the ecto-symbiotic association between free-living *Oophila* inside the embryonic egg capsule (Kerney, 2011). This intracapsular *Oophila* has a role in oxygenating the egg capsule microenvironment (Gilbert, 1942, 1944; Bachmann et al., 1986; Pinder and Friet, 1994; Mills and Barnhart, 1999; Bianchini et al., 2012) and potentially in removal of nitrogenous waste from the host (Goff and Stein, 1978; Bianchini et al., 2012; Small et al., 2014). Other modes of interaction between alga and embryo during the ecto-symbiotic stage of this association are relatively unexplored. One intriguing possibility is that the intracapsular *Oophila* fixes carbon from the atmosphere, uses energy from the sun to build fixed carbon into energy storage molecules like carbohydrates, and transfers that chemical energy to the salamander by exporting metabolically active compounds (Hammen and Hutchison, 1962; Goff and Stein, 1978; Graham et al., 2013, 2014). Such a mechanism is at play in other animal-alga photosymbioses, such as the coral-dinoflagellate mutualism, where the photosymbiont captures and transfers energy to the animal in the form of sugars and sugar alcohols like glucose and glycerol in nutrient-poor waters (Venn et al., 2008; Tremblay et al., 2012, Raven, 2017). However, none of these parallel naturally-occurring animal-algal photosymbioses include a vertebrate host.

There is an under-appreciated controversy in the published literature on metabolite transfer within the *Oophila*—*A. maculatum* symbiosis. Recent studies have found evidence of ^14^C-labeled photosynthate transfer from *Oophila* to their embryonic spotted-salamander hosts (Graham et al., 2013, 2014). However, an earlier study (Hammen and Hutchison, 1962), which also used a ^14^C label to detect intracapsular *Oophila* photosynthate transfer, came to the opposite conclusion: “The results of these experiments indicate that the facultative mutualism of *Ambystoma* embryos and the alga *Oophila* is not simply one of photosynthetic carbon dioxide fixation by the alga with subsequent transport of labeled carbohydrate to the embryo” (Hammen and Hutchison, 1962). Instead, this earlier report found that the embryos themselves were fixing a considerable amount of inorganic carbon, in the form of either carbon dioxide or bicarbonate ions. The possibility of heterotrophic carbon fixation was not explored by either recent paper and may be a confounding variable in their analysis and subsequent conclusions.

In aquatic systems, carbon dioxide (CO_2_) and bicarbonate (HCO_3_^-^) are critical for energy storage and central metabolism (Brinson et al., 1981; Boston et al., 1989). Heterotrophs primarily get the carbon they need from organic molecules like sugars and lipids while plants and algae collect carbon from the air or water and build it into the requisite organic compounds (Allen et al., 2005). There are, however, several metabolic processes common to both plants and animals that require inorganic carbon in the form of carbon dioxide or bicarbonate as substrates. Those include the urea cycle where the initial step of ammonia removal involves combining ammonium ions with bicarbonate and ATP by the enzyme carbamoyl phosphate synthetase to eventually remove the nitrogen in urea (Holden et al., 1999); the citric acid cycle where the enzyme pyruvate carboxylase combines carbon dioxide with pyruvate to produce oxaloacetate to fill in TCA intermediates (Jitrapakdee et al., 2006); and fatty acid biosynthesis where the first committed step involves adding bicarbonate to acetyl-CoA to produce malonyl-CoA, the fatty acid building block precursor (Blanchard and Waldrop, 1998).

Heterotrophic carbon fixation was first described by Harlan Wood in his studies of propionibacteria (Wood et al., 1941) and later in animals through his work on pigeon liver physiology (Wood et al., 1945; Kresge et al., 2005). The discovery of heterotrophic carbon fixation was at first met with skepticism, but later revealed in multiple bacterial and animal systems (Kresge et al., 2005). While there are six canonical autotrophic pathways to carbon fixation (Berg, 2011), many additional carboxylases have been characterized in heterotrophic cellular physiology (Erb, 2011). The scale of these “dark carbon fixation” pathways are often overlooked and may have considerable bearing on models of global carbon cycling (Baltar and Herndl, 2019). Coincidentally, many of the animal systems that revealed metazoan carbon fixation utilized amphibian embryos (Biggers and Bellve, 1974). Several definitive studies on carbon fixation by frog embryos were performed by Nobel laureate Stanley Cohen (Cohen, 1954, 1963), who later went on to discover nerve and epidermal growth factors (Shampo, 1999). Interestingly, the study of heterotrophic carbon fixation in amphibian embryos has received little research attention since the 1970’s despite decades of biochemical and molecular research into the mechanisms of development (Elinson and del Pino, 2011) and parallel research attention into the mechanisms of carbon fixation (Gong et al., 2016). In amphibians, carbon fixation was noted during early development of embryos of several frog species in the genus *Rana* (Cohen, 1954, 1963; Flickinger, 1954) and in the European newt *Triton* (Tiedemann and Tiedemann, 1954). Those early studies generally concluded that in early development carbon fixation proceeded via the action of carbamoyl phosphate synthetase, feeding into both de novo pyrimidine biosynthesis and nascent mRNA production, as well as directly into the urea cycle (Biggers and Bellve, 1974). Carbon fixation has been noted in other vertebrates as well, including fish (Mounib and Eisan, 1969) and mammals (Wales et al., 1969). In whole animals and embryos, the bulk of carbon fixation was observed to be tissue-specific reflecting the metabolic demands of different cell types (Biggers and Bellve, 1974).

Here, we revisit carbon fixation and translocation experiments taking care to separate algal and salamander contributions. We found that heterotrophic carbon fixation by salamander embryos is considerable and precludes our ability to distinguish metabolite transfer of algal fixed carbon from heterotrophic carbon fixation by the salamander using bulk measurements of whole embryos or egg capsules. Our results are incompatible with the notion that a measurable quantity of photosynthetically fixed carbon is transferred from alga to salamander in the egg capsule and are in agreement with the results from Hammen and Hutchinson, 1962. These results further indicate that more precise imaging of labeled carbon is required to definitively reveal whether algal metabolites are transferred to the host during the extracellular and intracellular portions of this symbiosis.

## Materials and Methods

### Egg mass collection and maintenance

*A. maculatum* egg clutches were collected from Michaux State Forest in central Pennsylvania (PA Fish and Boat Commission permit PA-727 type A) and Castamine Maine in March/April, 2019 (ME Department of Inland Fisheries and Wildlife permit 2020-590). *A. maculatum* egg clutches were maintained in modified Holtfreter’s solution (15 mM NaCl, 0.6 mM NaHCO3, 0.2 mM KCl, 0.2 mM CaCl2, 0.2 mM MgSO4•7H20) at 4°C to control the rate of development of the salamander embryos with constant light to maintain intracapsular algae. Prior to carbon fixation experiments, egg clutches were transferred to 18°C with a 12hr/12hr light/dark cycle for several days to facilitate development (Hammen and Hutchison, 1962; Graham et al., 2013, 2014). Development was monitored by visual inspection using a binocular microscope. When embryos reached Harrison stage 32-34 (Harrison, 1969) (Figure 1), individual eggs were removed from the jelly mass. Algal presence was confirmed by the green color of individual eggs.

**Figure 1.**
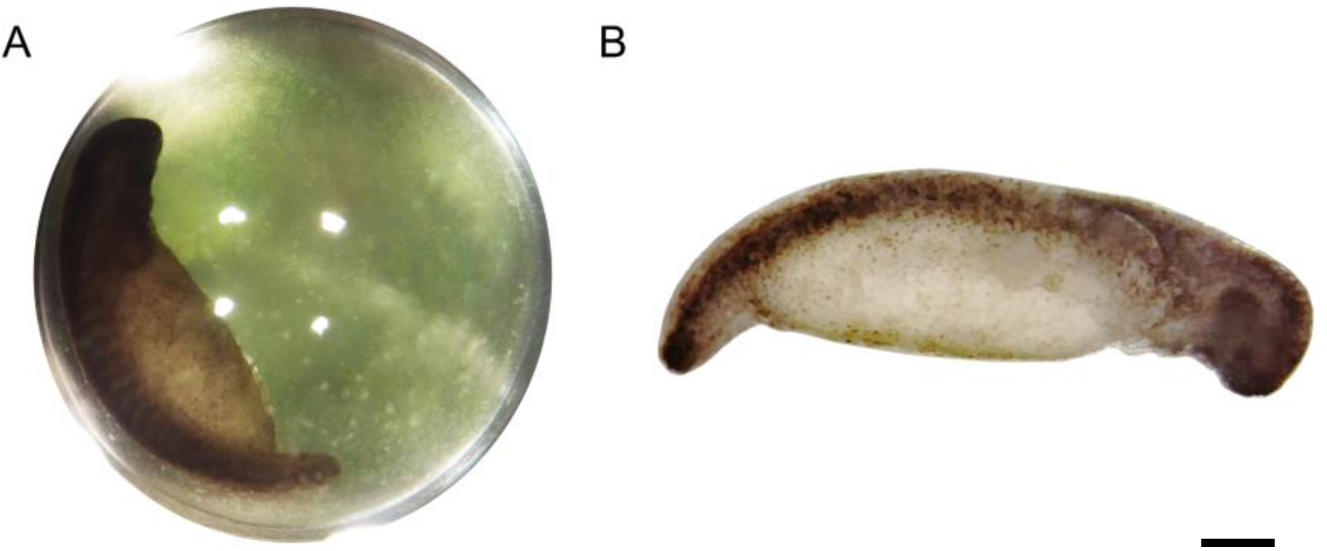
Typical stage 33 *A. maculatum* embryo from these experiments. **A.** Single egg capsule containing stage 33 embryo. Green hue is from intracapsular algae. **B.** Decapsulated stage 33 embryo. Scale bar: 1mm.

### ^14^C Bicarbonate incubation

#### Whole eggs

For experiments with whole eggs (Supplemental Figure 1), individual eggs were removed from the jelly mass by gentle manipulation with gloved hands, and transferred to individual scintillation vials in 2.5ml modified Holtfreter’s solution without sodium bicarbonate per egg. To each egg, 1.5 µl of ^14^C sodium bicarbonate [Perkin Elmer, 37 MBq/mL] was added. Adapting the protocol from Graham et al. (2014), 5 individual eggs per experiment were incubated in the light or dark for 2 hours. Following the initial incubation with ^14^C bicarbonate, eggs were removed from the solution and rinsed with DI water to remove unincorporated ^14^C bicarbonate from outside the egg. Following the rinse, eggs were transferred to 2.5ml modified Holtfreter’s solution (not radioactive) per egg and incubated in the dark for 2 additional hours. After the dark incubation, eggs were pierced with jeweler’s forceps, and the embryo was removed. Intracapsular fluid, including all algae, and the egg membrane were retained in one scintillation vial. Embryos were washed 3x with Holtfreter’s solution without sodium bicarbonate to remove carry-over algae and were placed into a separate scintillation vial. Embryos were homogenized in 500µl of 2M HCl and incubated for 24 hours in an open vial in a chemical fume hood to remove unincorporated sodium bicarbonate as CO_2_. 500µl of 2M HCl was added to the algal fraction, which was also incubated for 24 hours in an open vial in a chemical fume hood to remove unincorporated sodium bicarbonate. After 24 hours incubation, 10 mL of scintillation cocktail (Perkin Elmer, Ultima Gold LLT) was added to all scintillation vials before vortexing, and radioactivity was assayed using a Perkin Elmer Tri-Carb 3110TR scintillation counter. Results are reported as disintegrations per minute (DPM) as a measure of ^14^C incorporation.

### Decapsulated Embryo experiments

For experiments where embryos were removed from eggs prior to incubation with ^14^C bicarbonate, individual eggs were removed from the jelly mass by gentle manipulation with gloved hands. Eggs were pierced with jeweler’s forceps and embryos were transferred to clean Holtfreter’s solution without bicarbonate; algae were discarded. Embryos were transferred through clean Holtfreter’s solution without bicarbonate 3x to remove exogenous algae. Individual decapsulated and washed embryos were transferred to scintillation vials containing 2.5ml modified Holtfreter’s solution without bicarbonate. Controls consisting of killed embryos were completed by transferring decapsulated, washed embryos to a solution of 0.6x PBS containing 2.5% (w/v) glutaraldehyde and incubating the embryos in that solution for 15 minutes at room temperature. Following incubation in glutaraldehyde, embryos were washed 3x in Holtfreter’s solution without bicarbonate.

Labeled, cultured algae were made by adding 300 uL of ^14^C bicarbonate (at 37 MBq/mL) to 100 mL of *Oophila* algae in exponential growth in AF6 medium (at around 200,000 cells per mL) and incubating algae with ^14^C bicarbonate for 4 hours at 16°C under 260 µmol photons/m^2^/s. Following incubation, algae were transferred to 50mL falcon tubes and centrifuged at 300xg for 8 minutes. The supernatant was discarded and algae were resuspended in 20mL complete Holtfreter’s solution. Algae were pelleted a second time at 300xg for 8 minutes, and then resuspended in 1mL total Holtfreter’s solution. In parallel, unlabeled algae were subjected to the same centrifugation and wash procedure. Unlabeled algae from the resuspended pellet were counted on a hemocytometer.

For experiments with pre-labeled algae, a volume of ^14^C labeled algae approximately equivalent to 150,000 algal cells was added to live and glutaraldehyde killed decapsulated and washed embryos. Live and dead embryos were incubated with ^14^C bicarbonate (1.5 uL at 37 MBq/mL per embryo in 2.5mL Holtfreters without bicarbonate) or ^14^C labeled algae for 2 hours in the dark at 16 °C. Following incubation with ^14^C sodium bicarbonate or ^14^C labeled algae, embryos were washed 3x with unlabeled Holtfreter’s solution, transferred to individual scintillation vials, crushed in 500uL of 2M HCl, incubated for 24+ hours in open vials in a chemical fume hood to remove unincorporated bicarbonate, mixed with 10mL scintillation cocktail, and subjected to scintillation counting as above.

### Photosynthesis inhibitor experiments

Individual eggs were removed from the jelly mass by gentle manipulation with gloved hands. Six eggs per experiment were placed together in a 50 mL falcon tube in 15mL modified Holtfreter’s solution without sodium bicarbonate. Eggs were pre-incubated in the light at 16°C with two photosynthesis inhibitors, DCMU (3-(3,4-dichlorophenyl)-1,1-dimethylurea) and DBMIB (2,5-dibromo-3-methyl-6-isopropylbenzoquinone) at 20 µM final concentration for one hour. Following pre-incubation, 9 µl of ^14^C sodium bicarbonate, was added to each 50mL Falcon tube containing 6 eggs. Eggs were incubated in the light or dark for 2 hours. Following incubation, eggs were removed from the Falcon tubes, washed with DI water, opened with jeweler’s forceps, and separated into individual scintillation vials. Intracapsular fluid with algae and the egg membrane from one egg were collected in one vial, each embryo was washed 3x with Holtfreter’s solution and placed in a separate scintillation vial. Algae and embryos were processed as above for scintillation counting.

### Statistical analyses

Statistical analyses were performed in the R programming language (R Core Team, 2017). Plots were generated using the ggplot2 package (Wickham, 2016). One-way anova followed by the non-parametric Games-Howell post hoc test (used due to violation of homogeneity of variance in the data) was performed using the “userfriendlyscience” package (Peters, 2017).

## Results

### A. maculatum embryos fix carbon

Decapsulated *A. maculatum* embryos (Figure 1B) incubated in the dark with ^14^C bicarbonate incorporated inorganic carbon into their tissues as acid stable forms (Figure 2A) while control embryos without addition of ^14^C bicarbonate had no such signal (p<0.01). Decapsulated Embryos incubated in the dark exhibited significantly more ^14^C bicarbonate incorporation than total algae from whole eggs (as in Figure 1A) incubated in the dark (p<0.01) indicating that the elevated radioactive signal in the embryos could not have come from residual algae on or near the dark incubated embryos (Figure 2A).

**Figure 2.**
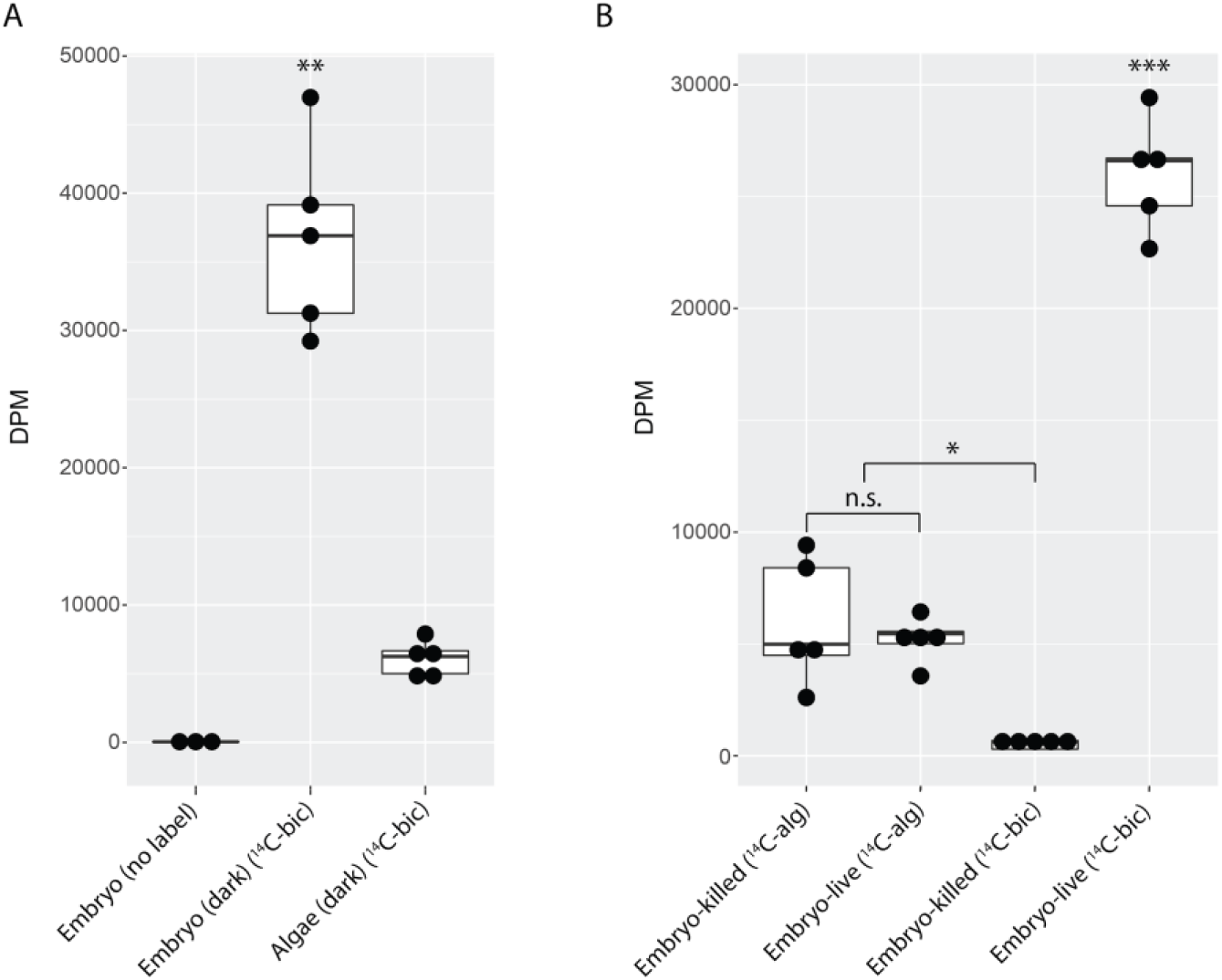
*A. maculatum* embryos fix carbon. Box and whisker plots showing raw data (dark circles) range (thin lines) averages (thick horizontal lines) and upper and lower quartiles (box). In each plot, the y-axis represents radioactivity assayed per sample (DPM, disintegrations per minute); the x-axis represents the treatments: **A.** embryo or algae incubated in the dark without (no label) or with ^14^C-bicarbonate (^14^C-bic); embryos incorporate significantly more carbon than algae in the dark. **B.** Glutaraldehyde-killed or live embryo (embryo-killed and embryo-live, respectively) incubated in the light with ^14^C-labeled alga (^14^C-alg) or with ^14^C-bicarbonate. Embryos must be alive for carbon fixation to take place. Pre-labeled algae do not impart additional benefits to live embryos compared to dead embryos. n.s.=not significant, *p<0.05, **p<0.01, ***p<0.001. Significance levels were determined by ANOVA followed by the Games-Howell post-hoc test.

To explicitly test whether residual algae that persisted through wash steps could account for the fixed carbon signal from embryos, *O. amblystomatis* cultures were incubated in the light with ^14^C bicarbonate for 5 hours, washed of free bicarbonate, and incubated in the dark with live or glutaraldehyde killed decapsulated and washed *A. maculatum* embryos. Such embryos exhibited an elevated fixed carbon signal compared to glutaraldehyde killed embryos incubated with ^14^C bicarbonate only (p<0.05), but there was no significant difference in acid-stable ^14^C measurements between live or dead embryos (p=0.9) (Figure 2B). The result suggests that some algae persist through the washes but living *A. maculatum* embryos do not actively accrue additional measurable photosynthate in these conditions. Decapsulated embryos incubated in the dark with ^14^C bicarbonate accumulated significantly more fixed carbon (Figure 2B) compared to embryos incubated with pre-labeled algae (p<0.001) or embryos killed with glutaraldehyde prior to incubation with ^14^C bicarbonate (p<0.001).

*A. maculatum* embryos compete with *O. amblystomatis* algae for bicarbonate Embryos from whole eggs incubated with ^14^C bicarbonate in the dark fixed significantly more carbon than embryos from whole eggs incubated in the light (Figure 3A, p<0.01). To test whether this was due to greater inorganic carbon availability when algae are not actively photosynthesizing (i.e. in the dark), photosynthesis inhibitors were added to the eggs in the light and the dark. Chemical inhibition of algal photosynthesis in the light decreased carbon fixation in the algae (Figure 3B, p<0.01), and resulted in elevated fixed carbon levels in salamander embryos, reproducing the effect of placing the egg in the dark (Figure 3C).

**Figure 3.**
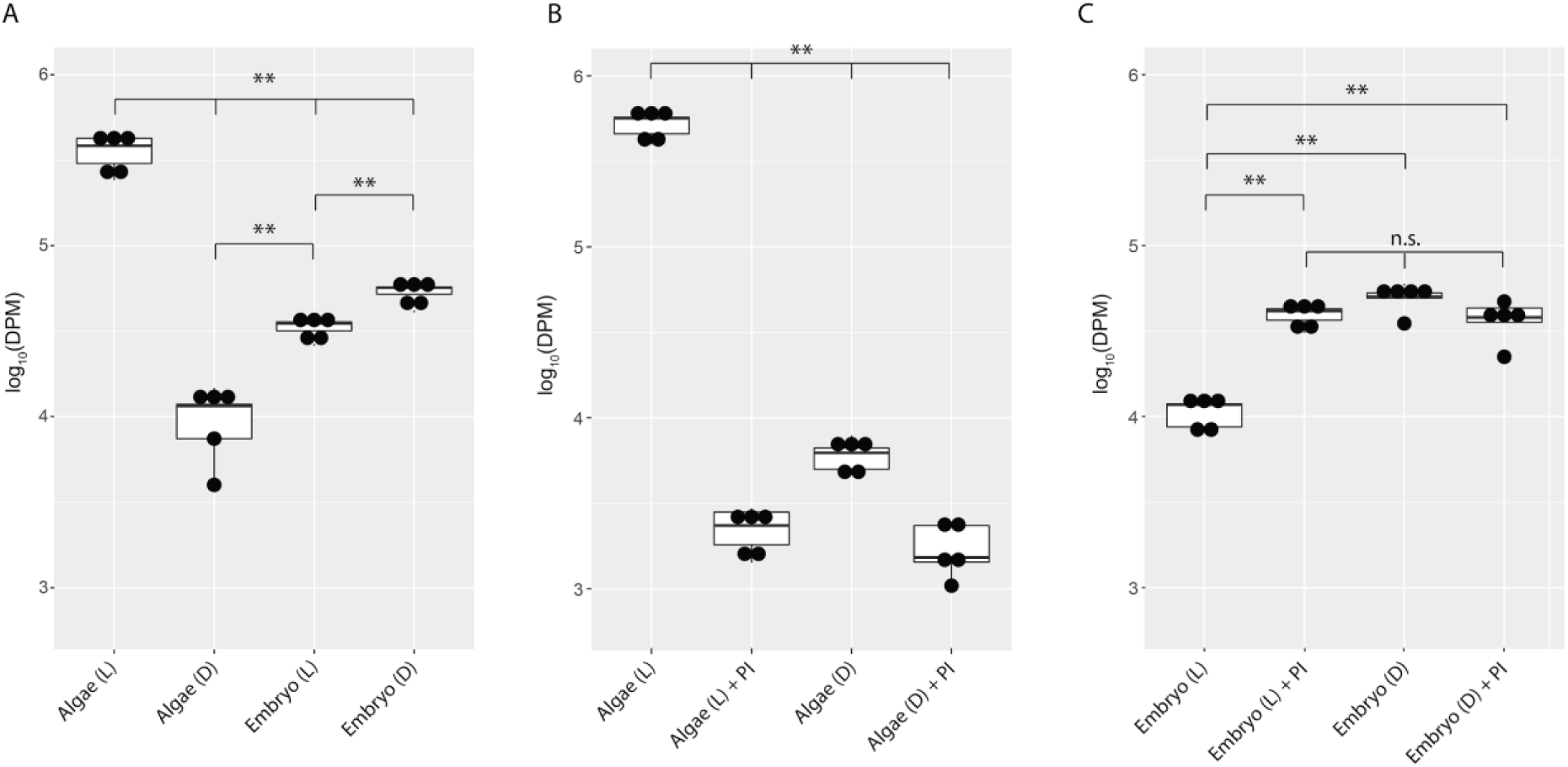
*A. maculatum* embryos compete with *O. amblystomatis* algae for bicarbonate. In each image, the y-axis represents radioactivity assayed per sample (DPM, disintegrations per minute); the x-axis represents the treatments. **A.** Algae from whole eggs incubated in the light (L) fixed significantly more carbon than algae in the dark (D) and embryos in the light or dark. Embryos in the light fixed more carbon than algae in the dark. Embryos in the dark fixed significantly more carbon than embryos in the light and algae in the dark. **B.** Photosynthesis inhibitors (PI) halt carbon fixation in algae in the light to a similar extent to keeping the algae in the dark. **C.** In the whole egg environment, photosynthesis inhibitors cause significantly increased carbon fixation in embryos, reproducing the effect seen when eggs are kept in the dark. n.s.=not significant, **p<0.01. Significance levels were determined by ANOVA followed by the Games-Howell post-hoc test.

## Discussion

In plants and algae, carbon dioxide is seen as a building block, a molecule captured from the air or surrounding fluid and fixed into organic matter as part of photosynthesis. In animals, carbon dioxide is largely a by-product of respiration that is also co-opted as an important biological buffer in body fluids. In addition to these roles, carbon dioxide and its hydration product, bicarbonate, can participate in several biosynthetic reactions outside of photosynthesis. One well studied and essential non-photosynthetic carbon fixation step that is present in animals and across cellular diversity is the direct conversion of pyruvate plus carbon dioxide to oxaloacetate by the mitochondrial enzyme pyruvate carboxylase (Sauer and Eikmanns, 2005; Marin-Valencia et al., 2010; Kim et al., 2016). Deficiency of pyruvate carboxylase in humans causes mortality during infancy (Marin-Valencia et al., 2010), indicating its critical importance to animals. Another important carbon fixation step in animals and across diversity is the first step in the urea cycle where bicarbonate, a nitrogenous compound (ammonia or glutamine), and ATP are combined to form the molecule carbamoyl phosphate (Allen et al., 2011). In humans, circulating citrulline, used in the inter-organ process of de-novo arginine biosynthesis (Curis et al., 2005; Breuillard et al., 2015; Marini et al., 2017), is largely synthesized from carbamoyl phosphate and ornithine in the intestines (Van De Poll et al., 2007). Given the quantity of citrulline (40 µM) in the blood of the average adult (Curis et al., 2005; Crenn et al., 2003), each person carries at least 2 mg of heterotrophically fixed carbon in their circulating blood. Carbamoyl phosphate is also used in de novo pyrimidine biosynthesis, a critical pathway in proliferating cells, assimilating inorganic carbon into nucleotides for use in a variety of downstream processes including DNA, RNA, and lipid synthesis (Fairbanks et al., 1995).

In terrestrial animals, the inorganic carbon incorporated in the above processes is best described as re-assimilated carbon. The bicarbonate and carbon dioxide reactants are derived from the end products of oxidative catabolism of organic molecules deposited in the blood, unlike in plants that pull carbon dioxide from the air. Aquatic animal eggs and embryos, however, have access to carbon dioxide dissolved in water, and can import that exogenous environmental carbon dioxide as bicarbonate for use in biosynthetic processes (Flickinger, 1954; Mounib and Eisan, 1973). In amphibian embryos and embryonic tissues, exogenous bicarbonate has been observed to enter the urea cycle in the liver (Brown Jr, 1962) and to get incorporated into a variety of compounds including pyrimidine nucleotides in whole embryos (Cohen, 1954; Flickinger, 1954).

Fixation of intracapsular carbon dioxide by amphibian embryos may be important for the localized pH regulation by specific cell populations. The bicarbonate buffering system is of critical importance to acid-base regulation in multiple physiological systems and is largely regulated by the dissociation of carbon dioxide in water, which lowers pH (Burggren and Bautista, 2019). Heterotropic fixation of carbon dioxide may help raise pH in specific tissue and cell populations. Amphibian embryos vary in their pH tolerances, and *A. maculatum* are particularly sensitive (Pierce, 1985). Their mortality can increase from <1% to >60% by lowering the pH from 7.0 to 6.0 and approaches 100% by pH 4.0 (Pough, 1976). Previous research in the *A. maculatum-Oophila* symbiosis has found a pH of 4.5 decreases the partial pressure of oxygen in the egg capsule and increases capsular ammonia as well as embryonic ammonia and lactate (Bianchini et al., 2012). These changes are inferred to be a direct effect on algal cellular physiology, which interferes with the net benefit *Oophila* confers to the host embryos. We have found no published evidence that a lower pH also increases rates of embryonic carbon fixation, which would be expected if these carboxylases are helping to regulate tissue-specific pH levels.

During the salamander-alga symbiosis, the same carbon fixation processes observed in other animals are likely active in *A. maculatum* embryos. The results presented here are consistent with heterotrophic fixation of exogenous bicarbonate by *A. maculatum* embryos. As demonstrated in other systems, such carbon fixation is necessary for proper development, and here we provide evidence that the algae and salamander compete for exogenously supplied bicarbonate for their various biosynthetic demands. Our results are consistent with the findings of Hammen and Hutchinson, 1962, suggesting that there is no measurable exchange of photosynthate from ecto-symbiotic algae to salamander embryos. The studies suggesting such an exchange (Graham et al., 2013, 2014) were likely only measuring variability in heterotrophic carbon fixation between individual embryos and did not control for the possibility that the salamander embryos themselves were fixing significant quantities of exogenously supplied bicarbonate.

Further studies on endosymbiotic algae are needed to reveal whether intracellular algal metabolites which are produced by the subset of endosymbiotic *Oophila* inside *A. maculatum* host cells are assimilated by their embryonic hosts. Our previous transcriptomic analysis has revealed the metabolic shift of intracellular algae from oxidative phosphorylation to fermentation (Burns et al., 2017), which may coincide with the formation of glycerol, formate or acetate (Catalanotti et al., 2013), which are used in other photosynthetic endosymbionts (glycerol; Le’on and Galva’n, 1995) or ectosymbiotic bacterial associations (formate and acetate; den Besten et al., 2013; Karasov and Douglas, 2013). While there is no evidence that endosymbiotic *Oophila* enable autotrophy of *A. maculatum* embryos, the chemical dialog between this intracellular mutalist and its vertebrate host is a fascinating research topic for subsequent studies.

## Acknowledgments

The authors thank the Challenge Yourself to Change (CYTC) community group in East Stroudsburg, PA, especially Rocky Sayles, Marquise Long, Dominic Kaps, and others who put on waders when collecting spotted salamander egg masses that contributed to this research. We thank Darryl Speicher, Roger Spotts, and Kettle Creek Environmental Education Center for aid in egg mass collection. We thank Hui Yang for thoughtful comments on drafts of the manuscript. The work was supported by Gordon and Betty Moore Foundation Grant #5604. The funders had no role in study design, data collection and analysis, decision to publish, or preparation of the manuscript.

## Author Contributions Statement

JB and SD conceived, planned, and conducted the experiments. JB performed data analysis. JB, SD, and RK discussed the results and implications and co-wrote the manuscript.

