## Supplemental Figure 1 for "Heterotrophic Carbon Fixation in a Salamander-Alga Symbiosis"

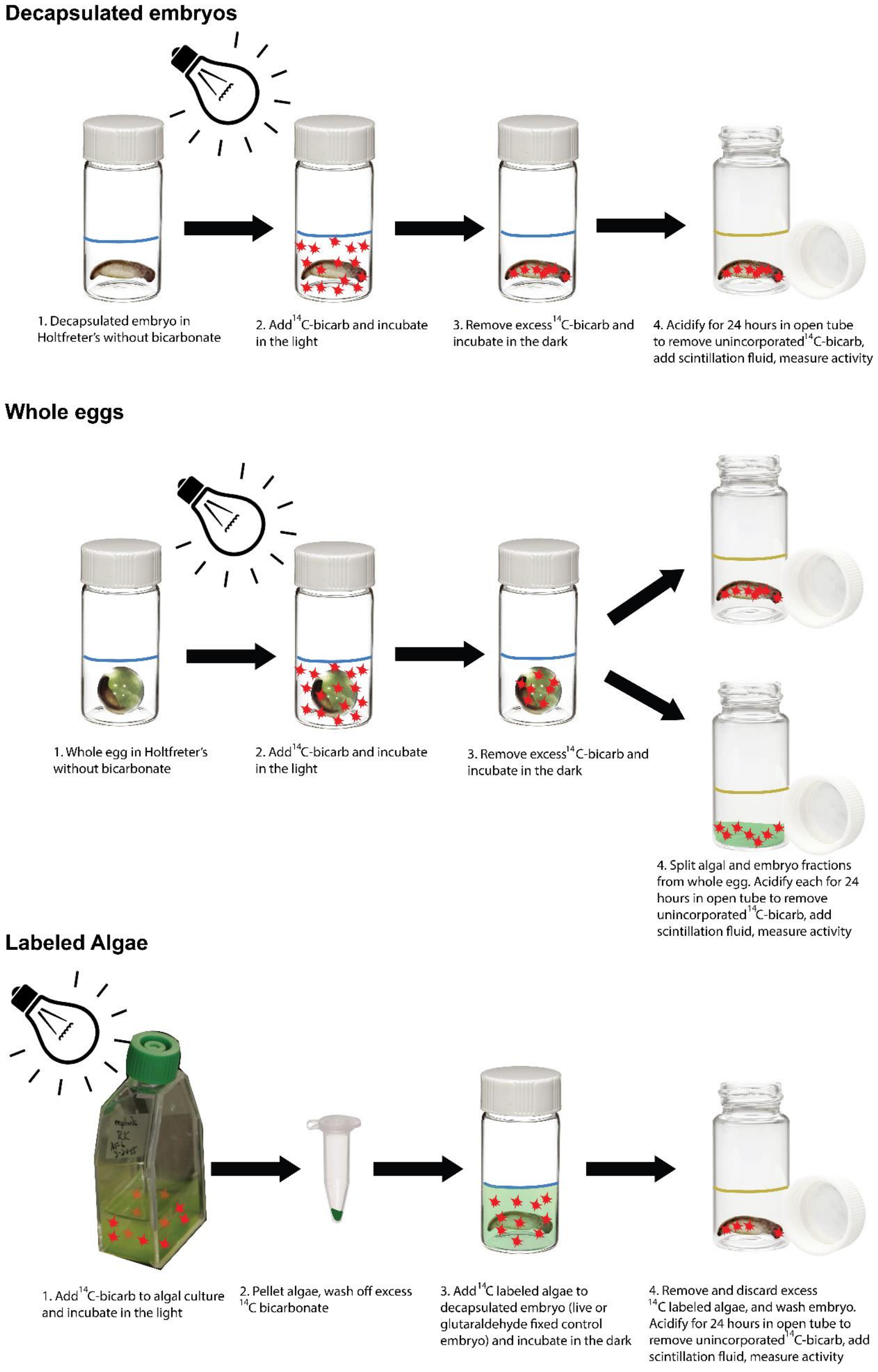
Decapsulated embryos
1. Decapsulated embryo in Holtfreter's without bicarbonate
2. Add $^{14}$C-bicarb and incubate in the light
3. Remove excess $^{14}$C-bicarb and incubate in the dark
4. Acidify for 24 hours in open tube to remove unincorporated $^{14}$C-bicarb, add scintillation fluid, measure activity
Whole eggs
1. Whole egg in Holtfreter's without bicarbonate
2. Add $^{14}$C-bicarb and incubate in the light
3. Remove excess $^{14}$C-bicarb and incubate in the dark
4. Split algal and embryo fractions from whole egg. Acidify each for 24 hours in open tube to remove unincorporated $^{14}$C-bicarb, add scintillation fluid, measure activity
Labeled Algae
1. Add $^{14}$C-bicarb to algal culture and incubate in the light
2. Pellet algae, wash off excess $^{14}$C bicarbonate
3. Add $^{14}$C labeled algae to decapsulated embryo (live or glutaraldehyde fixed control embryo) and incubate in the dark
4. Remove and discard excess $^{14}$C labeled algae, and wash embryo. Acidify for 24 hours in open tube to remove unincorporated $^{14}$C-bicarb, add scintillation fluid, measure activity

**Supplemental Figure 1. Basic experimental design**. **Decapsulated embryos:** (1) Decapsulated and washed embryos were placed in scintillation vials in Holtfreter's solution. (2) $^{14}C$-bicarbonate was added, and the specimens were incubated in the light for two hours. (3) The $^{14}C$-bicarbonate solution was removed and embryos were washed of excess $^{14}C$-bicarbonate, then transferred to the dark and incubated for 2 additional hours. (4) Decapsulated embryos were dissociated directly in HCl, acidified for 24 hours, scintillation fluid was added and samples were measured for activity. **Whole Eggs: (**1) Whole eggs were placed in scintillation vials in Holtfreter's solution. (2) $^{14}C$-bicarbonate was added, and the eggs were incubated in the light for two hours. (3) The $^{14}C$-bicarbonate solution was removed and intact eggs were washed of excess $^{14}C$-bicarbonate, then transferred to the dark and incubated for 2 additional hours. (4) Whole eggs were opened, and the algal and embryo fractions were separated. The embryo was washed 3x to remove carry over algae. HCl was added to the separate fractions, acidified for 24 hours, then scintillation fluid was added and samples were measured for activity. **Labeled algae: (**1) $^{14}C$-bicarbonate was added to cultured algae, which were incubated in the light for 4 hours. (2) Algae were pelleted and washed of excess $^{14}C$-bicarbonate. (3) $^{14}C$-labeled algae were added to live or dead decapsulated embryos and incubated in the dark for 2 hours. (4) Embryos were washed 3x to remove $^{14}C$-labeled algae, dissociated in HCl and acidified for 24 hours, then scintillation fluid was added and samples were measured for activity.
